# Predicting brain activity using Transformers

**DOI:** 10.1101/2023.08.02.551743

**Authors:** Hossein Adeli, Sun Minni, Nikolaus Kriegeskorte

## Abstract

The Algonauts challenge [Gifford et al., 2023] called on the community to provide novel solutions for predicting brain activity of humans viewing natural scenes. This report provides an overview and technical details of our submitted solution. We use a general transformer encoder-decoder model to map images to fMRI responses. The encoder model is a vision transformer trained using self-supervised methods (DINOv2). The decoder uses queries corresponding to different brain regions of interests (ROI) in different hemispheres to gather relevant information from the encoder output for predicting neural activity in each ROI. The output tokens from the decoder are then linearly mapped to the fMRI activity. The predictive success (challenge score: 63.5229, rank 2) suggests that features from self-supervised transformers may deserve consideration as models of human visual brain representations and shows the effectiveness of transformer mechanisms (self and cross-attention) to learn the mapping from features to brain responses. Code is available in this [github repository](https://github.com/Hosseinadeli/algonauts2023_transformers).

## 1 Introduction

A major goal of computational neuroscience is to model how different brain regions respond to visual input. An influential approach is to use image-computable models trained with supervised learning [Yamins and DiCarlo, 2016, Kriegeskorte, 2015] as a basis for linear encoding models [Kay et al., 2008, Naselaris et al., 2011, Gallant et al., 2012, St-Yves and Naselaris, 2018]. Linear encoding models, although arguably the simplest choice, are already high-dimensional (the number of parameters equals the product of the number of model units and the number of voxels to be predicted) and require strong regularization (L2 penalty equivalent to a 0-mean Gaussian prior on the weights) given the size of typical fMRI datasets. This has discouraged exploration of more complex mappings from model features to fMRI responses [Zhang et al., 2019, Ivanova et al., 2021].

However, the NSD 7-Tesla fMRI dataset [Allen et al., 2022] is more comprehensive than the datasets used in past studies. This dataset (and the Algonauts competition Gifford et al. [2023] based on it) give us the opportunity to explore other methods to map image features to neural activity. Our approach here leverages transformer neural networks for both the visual features and the mapping from features to voxels. The image features are extracted from a powerful transformer model trained with self-supervision and the mapping from features to fMRI voxels is learned using gradient descent. In post-processing, we take advantage of dimensionality reduction and regression methods to capture dependencies among fMRI responses within an ROI.

## 2 Related works

### Self-supervised Vision Transformers

Transformers have been shown to outperform convolutional neural networks (CNNs) on a variety of visual tasks including object recognition [Dosovitskiy et al., 2020]. In Transformers, the visual input is first divided into different patches and then encoded as feature vectors called tokens. At each layer of processing, a given token, that represents a particular image patch, updates its value by interacting with and mixing (“attending” to) the values of all other tokens that it finds relevant. The selective nature of this mixing has motivated naming this process “attention” in Transformers [Vaswani et al., 2017]. More recent studies have explored training these models on self-supervised objectives, yielding some intriguing object-centric properties that are not as prominent in the models trained for classification. When trained with self-distillation loss (DINO, Caron et al. [2021] and DINOv2 Oquab et al. [2023]), the attention values contain explicit information about the semantic segmentation of the foreground objects and their parts, reflecting that these models can capture object-centric representations without labels. Adeli et al. [2023] showed that the patch level feature representations from these models can be used to build models of attention spread to capture object-based attention and dynamics of object grouping in humans. These findings show that features from these models can be a good basis for predicting neural activity in the brain.

### Encoder-decoder Vision Transformers

Transformer-based encoder-decoder provide a general encoding and decoding framework that has achieved great performance in many domains [Vaswani et al., 2017] including domain where one modality (e.g. image) is mapped onto another one (e.g. language). This framework has been recently applied to the problem of object detection and grouping in images (DETR; Carion et al. [2020]) yielding state-of-the-art performance. The encoder in this model converts the image to rich object-centric features. The decoder uses learnable embeddings, called queries, corresponding to different potential objects, that gather information from the encoder features using cross-attention over several layers. Different object queries also attend to one another using self-attention in the decoder. After the decoding process, each object query can then be linearly mapped into to the category and bounding box for an object. The model is trained end-to-end and can detect many objects in one feedforward pass.

### Predicting primate visual representations with self-supervised neural network models

Recent work has shown that convolutional and recurrent neural networks trained using self-supervised contrastive losses (such as SimCLR; Chen et al. [2020]) match the predictive power of supervised models for high-level ventral-stream visual representations in the brain [Konkle and Alvarez, 2022, Chen et al., 2022]. These works argue for the self-supervised learning methods as a more plausible objective function for learning brain like visual representations.

## 3 Dataset and Challenge

The Algonauts 2023 challenge makes use of the NSD dataset Allen et al. [2022]. Briefly, the fMRI responses were collected from 8 subjects, each seeing up to 10000 images. The images were presented three times each and the average responses are provided as part of the challenge dataset. This dataset also focuses on a subset of ROIs that are more visually responsive. The last three sessions for each subject are held out as test set. The training set for each subject provides fMRI response for about 19k and 20k vertices for the Left and the Right Hemispheres (LH and RH), respectively, in response to each image. The behavioral data are also provided as part of the NSD dataset for the all the experimental trials along with single trial level fMRI responses. Also provided are ROI and stream level labels for each vertex. The ROI level labels are provided for early visual areas (‘V1v’, ‘V1d’, ‘V2v’, ‘V2d’, ‘V3v’, ‘V3d’, and ‘hV4’), body selective areas (‘EBA’, ‘FBA-1’, ‘FBA-2’, and ‘mTL-bodies’), face selective areas (‘OFA’, ‘FFA-1’, ‘FFA-2’, ‘mTL-faces’, and ‘aTL-faces’), place selective areas (‘OPA’, ‘PPA’, ‘RSC’), and word selective areas (‘OWFA’, ‘VWFA-1’, ‘VWFA-2’, ‘mfs-words’, and’mTL-words’). The stream level mapping assigns vertices to different streams (‘early’, ‘midventral’, ‘midlateral’, ‘midparietal’, ‘ventral’, ‘lateral’, and ‘parietal’ and ‘unknown’). The ‘unknown’ label is used for all the vertices that were not assigned to an ROI or a stream in each of these mappings.

## 4 Approach

We apply the the general transformer encoder-decoder model to map images to fMRI responses. Figure 1 shows the architecture for our model. The input image is first divided to 31× 31 patches of size 14 × 14 pixels. These image patches are input to the encoder model which is a 12-layer vision transformer (with token size of 768) that is trained using the DINOv2 self-supervised method [Oquab et al., 2023]. The encoder is frozen after pre-training and is used as a backbone to provide features. These features can be extracted from any layer of the encoder as they all have the same dimensionality (31× 31× 768). Given the token size of 768, it would not be feasible for our decoder to input features from all the layers as it would signifcalty increase the number of trainable parameters in our model. In transformers, each token is represented with three vectors: key, query, and value. Any of these vectors could be used as patch level representations. We selected query features as they have been shown to have a more object-centric feature representations Adeli et al. [2023].

**Figure 1:**
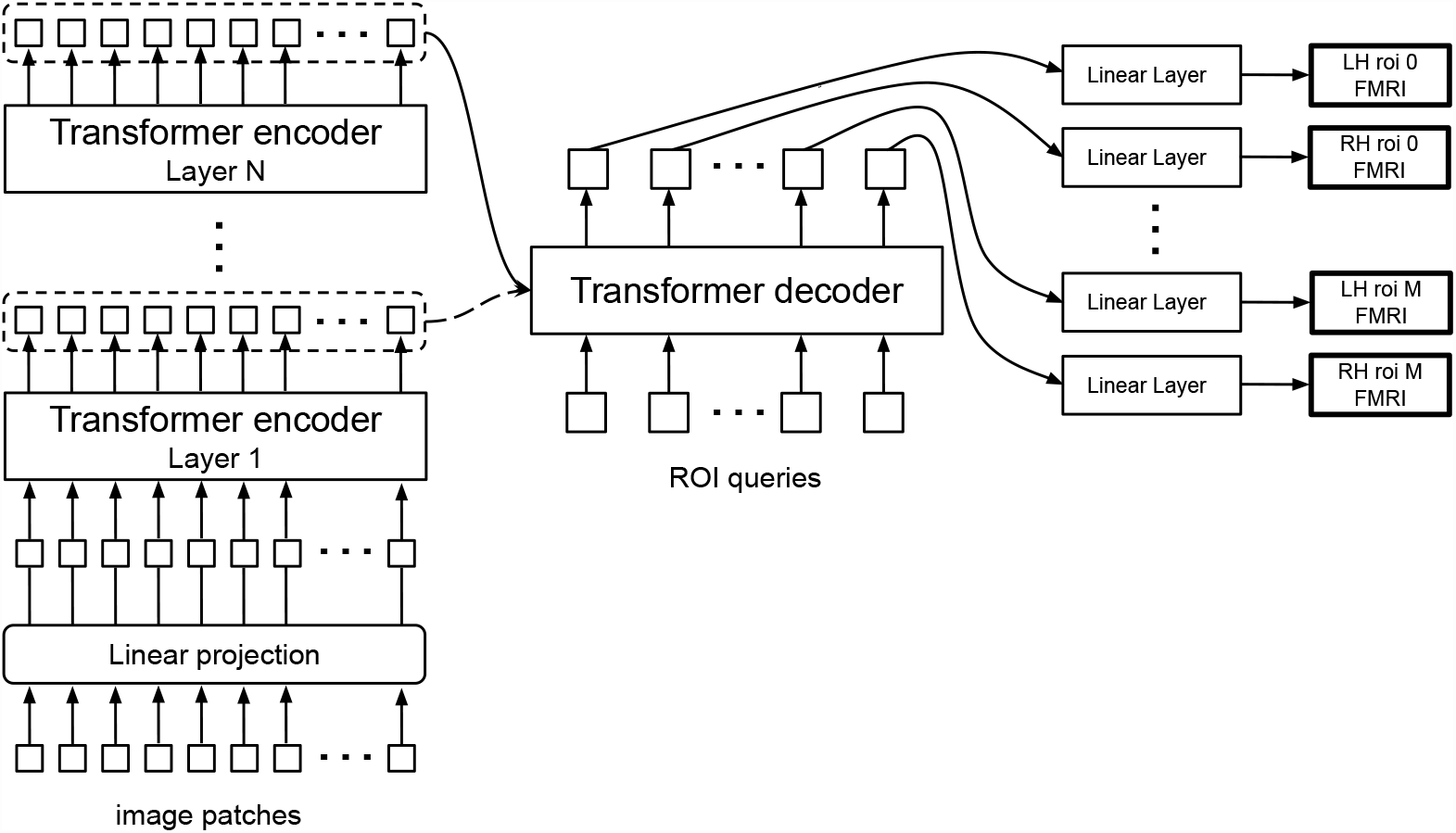
Model Architecture

The decoder uses input queries corresponding to different brain ROIs in different hemispheres to gather relevant information from the encoder output for predicting neural activity in each ROI (in the manner of Carion et al. [2020]). These ROI queries can also attend to one another through self-attention. Note that these queries are learnable embeddings and are different from the feature queries that area linear mapping of each token representation. We use a single-layer transformer for the decoder with one cross-attention and one self-attention operation. The output ROI queries are then mapped using a single linear layer to fMRI responses of the corresponding ROI. In our implementation, each ROI token is linearly mapped to a vector with the size equal to the number of vertices in that hemisphere. The response is then multiplied by a mask that is zero everywhere except for the vertices belonging to that ROI. This masking operation ensures that the gradient signal feeding back from the loss will only train linear mappings to the vertices of the queried ROI. The responses from different ROI readouts will then be combined using the same masks to generate the prediction for each hemisphere. The decoder and the linear mappings are trained with the Adam optimizer [Kingma and Ba, 2014] using mean-squared-error loss between the prediction and the ground truth for each image. We train the models separately for each subject.

### 4.1 Model experiments

We train models with different combinations of the encoder output layer and decoder queries as we anticipate different ROIs would be better predicted using visual input from different levels of abstraction in the encoder and different levels of granularity in decoder mapping.

When decoder tokens correspond to the different streams, the model can provide predictions for all the vertices. In this case, we use 16 decoder queries, 8 for each hemisphere (7 streams + one for all the vertices labeled ‘unknown’). For each subject we create a random 90/10 split for training and validation. The trained model can then be tested on the validation set to examine its performance. Figure 2 shows ROI specific, stream specific and overall correlations between the model predictions and the ground truth for one validation set. In this model the query features from the last layer of the encoder are used as the input representation to the decoder. We number the layers starting from the last layer of the encoder with the layer index increasing to 12 when we reach the earliest layer in the encoder, therefore in this case the input layer would be layer 1.

**Figure 2:**
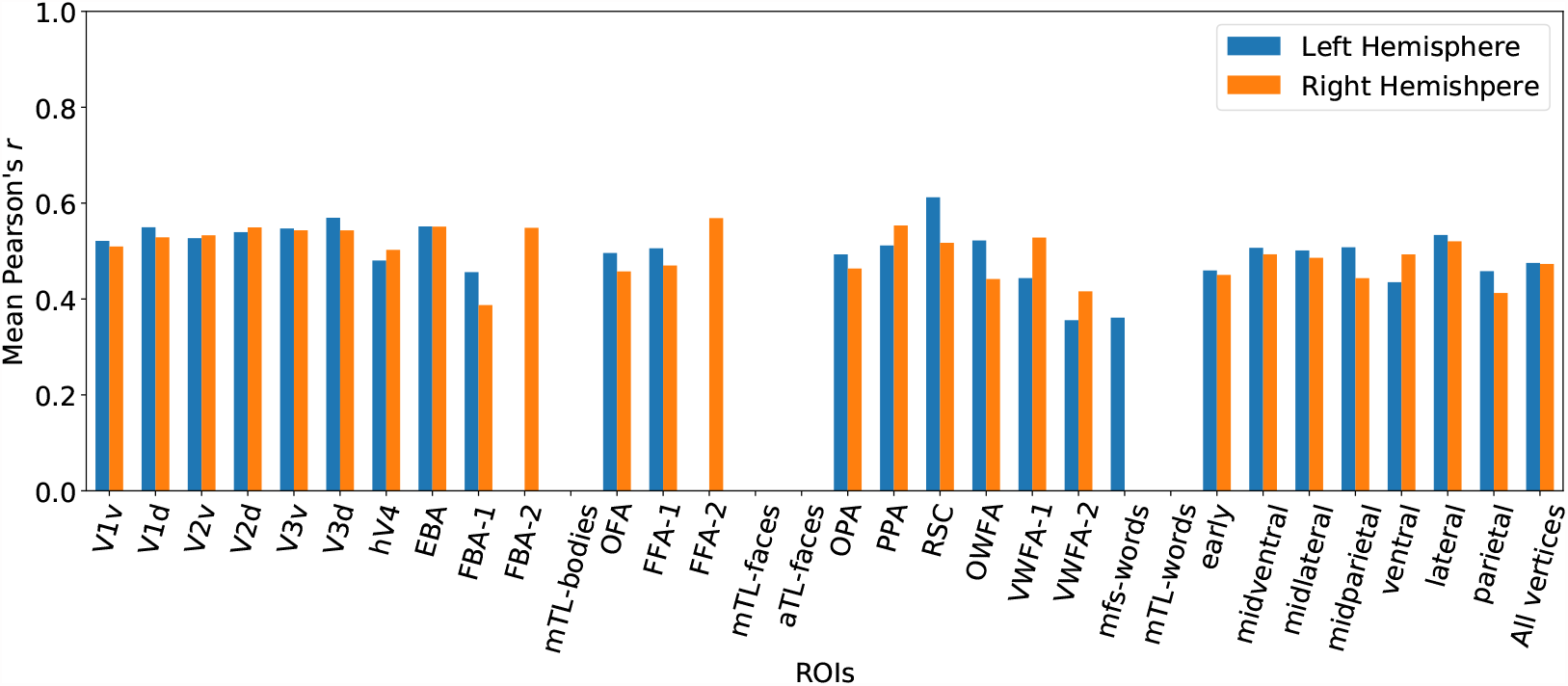
ROI- and stream-specific model prediction performance for a stream decoder for subject 1

Figure 3 shows the correlations for a different model, where the input layer is 8 and the decoder queries are based on the early visual areas. In this case the model does not predict anything for all the vertices with the ‘unknown’ tag so the predictions are focused on the ROIs. Comparing this figure with Figure 2 shows that the predictions improve for early visual areas.

**Figure 3:**
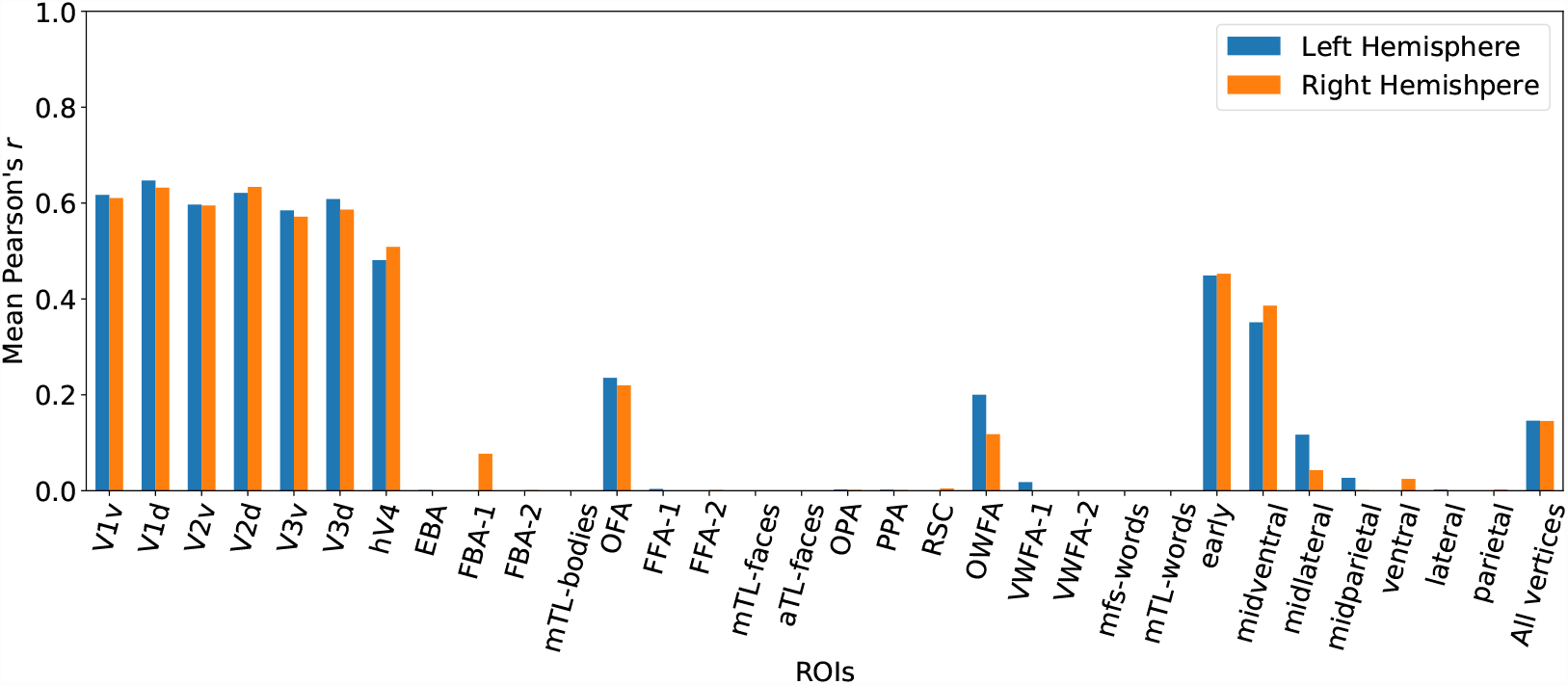
ROI- and stream-specific model prediction performance for a visual decoder for subject 1

### 4.2 Postprocess and ensemble

Based on some initial piloting, we settled on 22 models using different combinations of encoder output layer and the decoder ROIs (Table 1). We perform 10 fold Cross validation for each of our 22 models and use a different initialization seed for each fold. This process yields model predictions for all the images in the training set for each subject along with initial predictions for the test split.

**Table 1:** All model run specifications

| encoder output layer<br>(1 corresponds to the top layer) | decoder ROIs |
| --- | --- |
| 1, 2, 3, 4, 5, 6 | ('early', 'midventral', 'midlateral', 'midparietal', 'ventral', 'lateral', and 'parietal' and 'unknown') |
| 5, 6, 7, 8, 9 | ('V1v', 'V1d', 'V2v', 'V2d', 'V3v', 'V3d', and 'hV4') |
| 1, 2, 3, 4 | ('EBA', 'FBA-1', 'FBA-2', and 'mTL-bodies') |
| 4 | ('OFA', 'FFA-1', 'FFA-2', 'mTL-faces', and 'aTL-faces') |
| 1, 2, 3, 4 | ('OPA', 'PPA', 'RSC') |
| 3, 4 | ('OWFA', 'VWFA-1', 'VWFA-2', 'mfs-words', and 'mTL-words') |

With these initial predictions of the neural activity for the the training set, we can now apply dimensionality reduction techniques and add other features to the mix. For each subject we apply PCA to the model predictions for all the training images to get 100 pcs for each hemi-sphere. We then concatenate the reduced model predictions with 14 behavioral variables for each given image. These variable are: [‘RUN’, ‘TRIAL’, ‘TIME’, ‘ISOLD’, ‘ISCORRECT’, ‘RT’, ‘CHANGEMIND’, ‘MEMORYRECENT’, ‘MEMORYFIRST’, ‘ISOLDCURRENT’, ‘ISCORRECTCURRENT’, ‘TOTAL1’, ‘TOTAL2’, ‘BUTTON’] (detailed explanation of the variables see https://cvnlab.slite.page/p/fRv4lz5V2F/Behavioral-data). Each image was presented three times and we naively average these values for the three presentations. We then train linear regression to map from the model predictions + behavioral features to 100 PCs of the fMRI signal, from which we can reconstruct the fMRI predictions. The benefit of this approach over directly predicting the fMRI signal is that it captures the dependencies between the vertices.

We use 10-fold cross validation again to learn this second mapping. This postprocess can be applied to all the model predictions to yield new predictions of the test set and the validation set which can be evaluated to provide a goodness of the prediction for all vertices. Averaging all the runs for a given model setting, we can get one prediction for test set from that model and one vector determining the goodness of the model prediction for all vertices based on average performances on the validation set in each split.

In the final stage, for each vertex we combine the values across all the models using a softmax operation. The softmax weights are based on goodness of the prediction for each model for that vertex. Our model achieves a score of 63.5229 on the test set and our code is available in this [github repository](https://github.com/Hosseinadeli/algonauts2023_transformers).

